# Minimizing the number of phosphenes required for object recognition under prosthetic vision

**DOI:** 10.1101/2024.09.26.614440

**Authors:** Elsa Scialom, Udo A. Ernst, David Rotermund, Michael H. Herzog

**Author notes:** **Corresponding author:** Elsa Scialom. **Funding information:** All authors were funded by ERA-NET Neuron I-See “Improving intracortical visual prostheses using complex coding and spontaneous activation states” (32NE30_198552; BMBF 01EW2104A). U.A.E and D.R received additional funding from the Stiftung Bremer Wertpapierboerse. **Commercial relationships:** None.

## Abstract

Cortical prostheses offer the potential for partial vision restoration in individuals with blindness by stimulating V1 neurons to produce phosphenes. However, the low number of phosphenes that can be elicited in practice makes encoding of whole objects difficult, and the round shape of phosphenes lack the contour cues necessary for perceptual grouping. We propose a minimalistic encoding approach that focuses on essential visual information. We fragmented objects’ contours into either phosphenes or curved segments, providing either low or high local visual information. 46 participants identified these fragmented objects in a free-naming task. The number of fragments gradually increased to quantify the minimum number of phosphenes and segments necessary to recognize objects. Most objects could be recognized with only 65 phosphenes, which is in the range of implantable electrodes in human patients. Participants required 27% fewer segments than phosphenes to recognize objects. Including individual objects as a random effect in a linear mixed model substantially increased the explained variance, suggesting that the minimal number of fragments required for object recognition in prosthetic vision strongly depends on the particular object. Our results demonstrate that a minimalistic approach can substantially reduce the number of phosphenes required for recognition, emphasizing the importance of identifying critical object features to minimize brain stimulation in visual prostheses.

## Introduction

Around 40 million people worldwide are blind. Blindness is associated with physical and mental comorbidities dramatically affecting the life quality of blind patients (Court et al., 2014). Inspired by the success of cochlear implants, several research initiatives were set up to develop visual prostheses (for a review see Borda & Ghezzi, 2022). Initially, research focused mainly on retinal prostheses (e.g., Luo & Da Cruz, 2016; Ferlauto et al., 2018). Since patients with advanced retinitis pigmentosa or optic nerve damage cannot benefit from retinal prostheses, cortical prostheses were developed.

Today’s cortical prostheses aim at providing visual percepts to the blind by stimulating neurons of the primary visual cortex (V1). They consist of a camera capturing images of the blind person’s surroundings which are transmitted to a processor. The processor converts the signal via a stimulator into electrical impulses which are then transmitted into the brain tissue by an implant in V1 (Shepherd et al., 2013). Such stimulation of the cortical tissue elicits discrete percepts of light patches called “phosphenes”. Currently, V1 prostheses could elicit only 88 phosphenes in human patients implanted with an Utah array of 10×10 electrodes (Fernández et al., 2021). However, this resolution is not sufficient to enable object recognition, as shown by simulated prosthetic vision studies indicating that the threshold for object recognition is somewhere between 256 and 576 phosphenes (Zhao et al., 2010). Hence, simulated prosthetic vision studies with healthy participants are central to identify how the world should be rendered under prosthetic vision (Pio-Lopez et al., 2021).

Current studies use various ways to preprocess the image captured by the camera of the prosthesis (for a review see Wang et al., 2022). The most common preprocessing algorithm is to use edge extraction of the input image (e.g., Zhao et al., 2010; Thorn et al., 2020). Recently, more sophisticated optimization techniques have been used. For example saliency-based segmentation in which the most salient parts of the captured scenes are rendered (e.g., Li et al., 2018), as well as deep-learning techniques to simplify the layout of a scene (e.g. Sanchez-Garcia et al., 2020). The latter studies show better navigation and/or object recognition performance with preprocessed images.

These studies typically simulate simultaneous stimulations of all electrodes present along object contours. However, such stimulation of numerous electrodes can be detrimental in two ways. First, stimulating electrodes of cortical implants lead to increased tissue damage compared to passive electrodes which are implanted but not stimulating (Rosenfeld et al., 2020). Second, the simultaneous perception of hundreds of phosphenes makes perceptual grouping difficult for patients. Indeed, hundreds of phosphenes are still too few to determine which phosphenes belong to the same object and which do not. This problem is further enhanced by the isotropic perceptual nature of phosphenes, as their round appearance lacks contour cues that e.g., oriented segments could provide.

To address these limitations, we adopt a minimalistic approach hypothesizing that participants will be able to recognize objects even with very few phosphenes placed around the object’s contours. Second, we hypothesized that if observers use the directional local visual information carried by curved segments, they will need less segments to recognize objects compared to roundish phosphenes providing isotropic local visual information. To test these hypotheses, we use a traditional psychophysical study design which positions fragments along the contours of an object (as in e.g., Panis et al., 2008; Elder et al., 2018) and gradually increase the number of fragments to quantify the minimum number of fragments necessary for object recognition. To vary the local visual information provided by fragments, we fragmented objects using an algorithm placing either phosphenes or curved segments along the object’s contours, respectively providing low and high local visual information.

## Methods

### 1. Participants

We recruited 63 healthy volunteers from the Ecole Polytechnique Fédérale de Lausanne (EPFL) and the University of Lausanne. Fourteen participants undertook a labelling task to provide ground truth for an object recognition task performed by a new set of 49 participants. Since both the labelling and the object recognition tasks were free naming tasks, we only included native French speakers. One participant from the labelling task and three from the object recognition task were excluded because of insufficient French. Thus, data from 13 participants (*n*_males_ = 7, *M*_age_ = 21.5, *SD =* 1.6) for the labelling task and 46 participants for the object recognition task (*n*_males_ = 33, *M*_age_ = 21.4, *SD* = 2.6) were considered for the respective data analyses. All participants had normal or corrected-to-normal vision, assessed by the Freiburg Visual Acuity test (Bach, 1996). Participants provided written consent for their participation and were compensated with 25 Swiss francs per hour. The study was approved by the local ethics committee of the canton of Vaud in Switzerland (CER-VD).

### 2. Apparatus

The participants were seated in a dimly illuminated room (0.5 lux), 50 cm away from the screen in the labelling task and 150 cm away in the object recognition task. Stimuli were presented on an ASUS VG248QE LCD monitor (53×30 cm, 24”) with a screen resolution of 1920×1080 pixels and a refresh rate of 120 Hz. The maximum luminance of the screen was 100 cd/m^2^. The white point of the monitor was adjusted to D65. The monitor was calibrated for a transfer function with gamma=2.2.

The experiment runs under Psychtoolbox (Brainard, 1997) and MATLAB (MathWorks Inc., Natick, MA, USA).

### 3. Stimulus generation

We selected a subset of images from the Bank of Standardized Stimuli (BOSS; Brodeur et al., 2010), depicting everyday objects. These objects were then fragmented either with phosphenes or with curved segments. We fragmented objects with an algorithm that takes an RGB(A) image as input, extracts contours, fragments the contours, and then reduces fragment density (see Figure 1). Figure 2 shows example objects fragmented with our algorithm.

**Figure 1.**
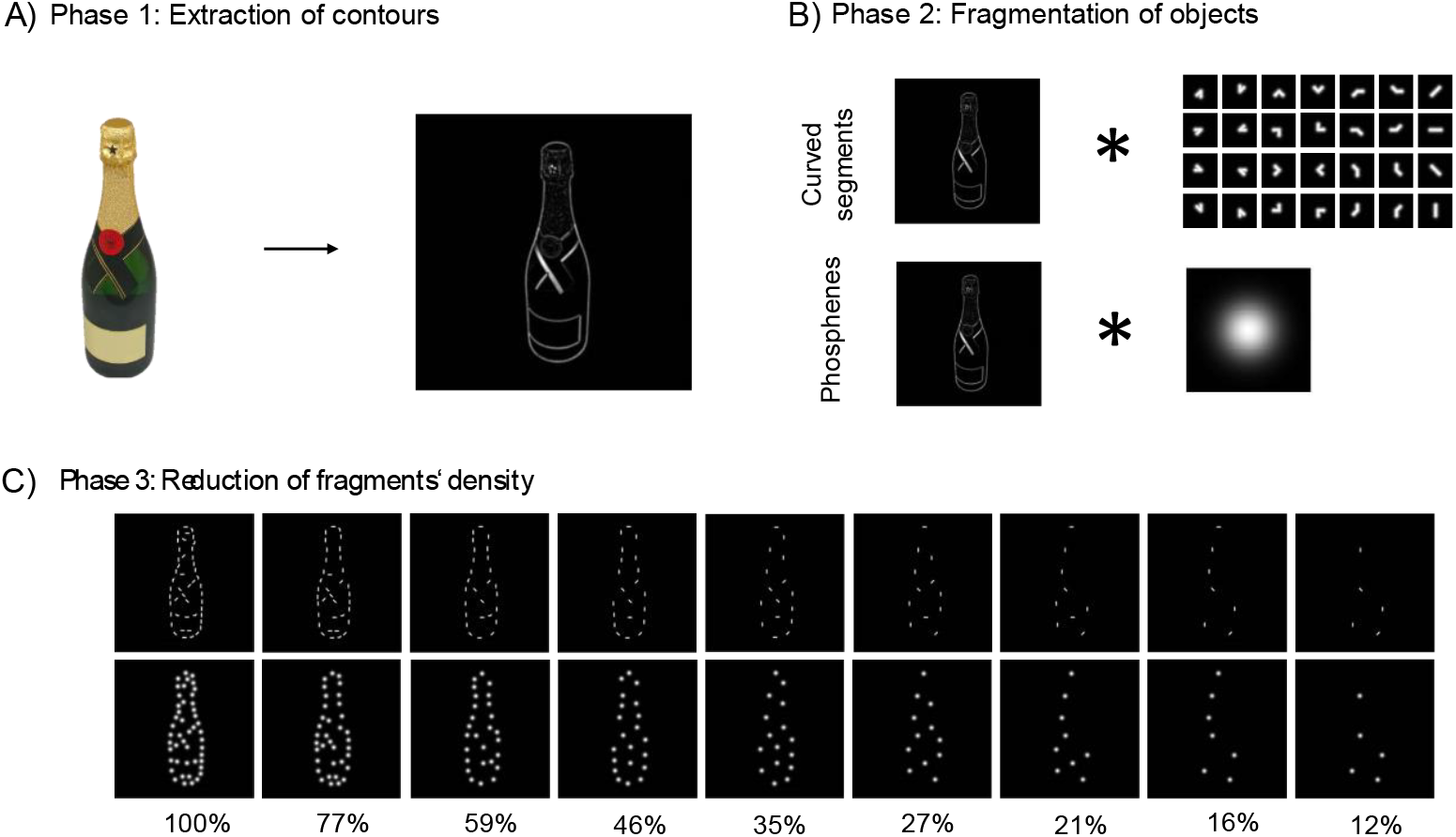
Algorithmic fragmentation of objects. A) Contours of images from the Bank of standardized stimuli (Brodeur et al., 2010) are extracted. B) Contours are fragmented by placing curved segments at positions where they have a large overlap with the object’s contours. This is done with a convolution and summation. Phosphenes are placed at the same pixel coordinates as the segments. C) Fragments’ density is reduced such that fragments with the least overlap were removed first until 77%, 59%, 46%, 35%, 27%, 21%, 16%, and 12% of fragment’s densities were reached. All colored images were taken from Brodeur et al., (2010) and are licensed CC-BY-NC-SA.

**Figure 2.**
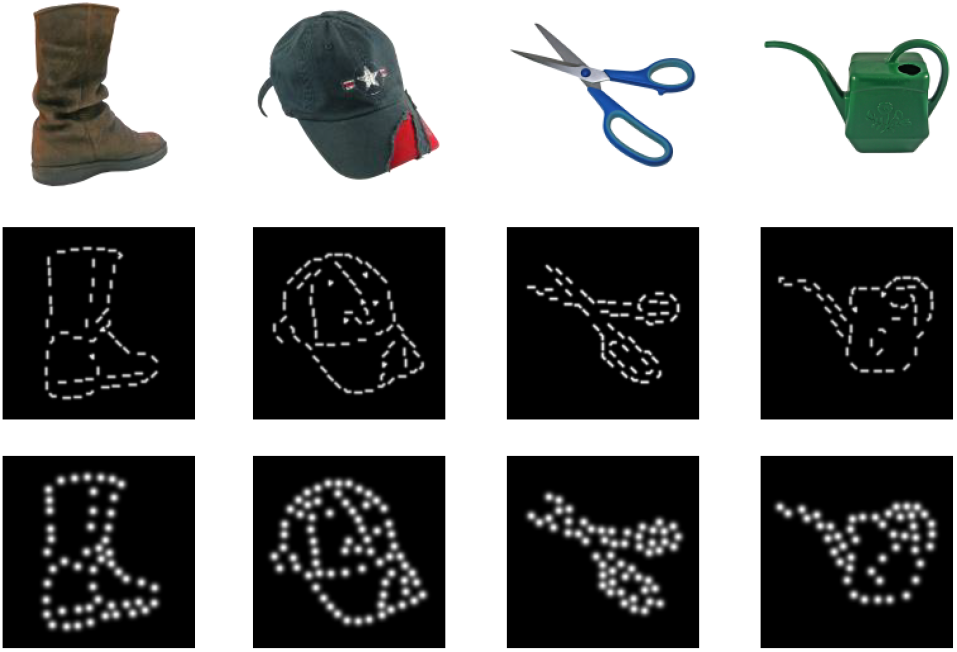
Example stimuli obtained with our algorithm fragmenting objects. Top row shows the input images, middle row shows the rendered images as curved segments (100% fragment density) and the bottom row shows images as phosphenes (100% fragment density). Colored images were taken from Brodeur et al., (2010) and are licensed CC-BY-NC-SA.

In the following, we summarize the procedure and refer readers for details of the implementation to: https://github.com/davrot/percept_simulator_2023/blob/main/offline_encoding.py

#### a) Contour extraction

The BOSS contains objects displayed on a white background. Since some images contained pixels marked to be transparent, we first converted these pixels to white before transforming the RGB representation to gray scale. To avoid boundary effects during contour extraction and fragmentation, the images were padded with a sufficiently large border having the same background color, and resized so that each object in its fragmented representation would extend over roughly 10° of visual angle (either horizontally or vertically). For contour extraction, we used a standard Gabor filter bank with 8 orientations, containing sine- and cosine-templates forming quadrature pairs allowing phase-invariant contour extraction.

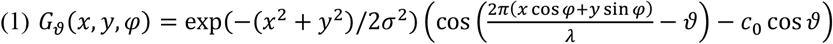

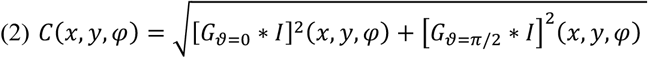

With *c*_0_ = exp(−2(*πσ*/*λ*)^2^), the star * denoting convolution (via a two-dimensional Fast Fourier Transform), and *I*(*x, y*) being the gray-level image. The result *C*(*x, y, φ*) indicates the strength at which a local contour of orientation *φ* is present at image location (*x, y*). Parameters for filter size *σ* = 0.06 arcdeg and spatial scale *λ* = 0.12 arcdeg were kept constant.

#### b) Object fragmentation

For fragments, we used two representations, one containing a single Gaussian shape with width *σ*_*G*_ = 0.18 arcdeg (a “phosphene”), and one containing a set of “clock hands” (Mineault et al., 2013) pointing into different directions with variable intermediate angles (“curved segments”, see below for detailed description and Figure 1B). Each fragment *i* was represented in form of a filter 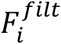 (*x, y, φ*) with explicit representation of orientation for the *fragmentation process*, and in form of an image patch 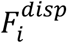 (*x, y, φ*).

Fragmentation started by first computing the overlap *O*_*i*_(*x, y*) of any curved segment fragment to any location in the contour representation *C*(*x, y, φ*) by means of a convolution and summation over the Gabor’s orientation variable *φ*:

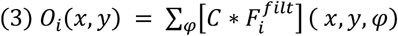

Locations (*x*’, *y*’) and identity *i*’ for best-matching fragments were then determined by an iterative procedure which selected the best match from *O* multiplied by a mask *M*(to keep the segments from overlapping) of candidate positions still eligible for placement of a fragment:

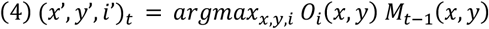

Here, *t* denotes iteration step with *M*_0_(*x, y*) = 1 being the initial state of *M*. After each selection, the mask of candidate positions was updated by excluding all positions within a radius *r*_*excl*_ = 0.65 arcdeg around the selected position (*x*’, *y*’)_*t*_, and by attenuating the overlap *O* beyond that radius with a factor proportional to a Gaussian with width *σ*_*disc*_ = 1.2 arcdeg centered at (*x*’, *y*’)_*t*_, according to the following equation where *θ* denotes the Heavyside function and 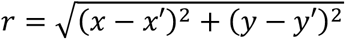:

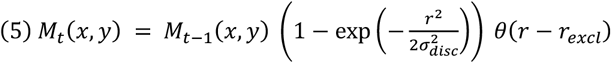

We stopped after placing *N*_*ele*_ = 100 fragments, which provided a sufficiently dense representation. Since the algorithm tends to place fragments at positions with uninformative image content (i.e. low-amplitude pixel noise at otherwise homogeneous image patches) when locations at or around objects become exhausted, we selected only those locations at which the overlap *O* exceeded a value of 10% of the maximum overlap. For allowing to display a suitable range of different fragment densities for the same object, we required the remaining number of fragments *N*_*rem*_ to be at or above 40, otherwise the image was not used in our study. The output of the fragmentation step consisted of a list of the type of fragments, their positions, and their quality of overlap with the contour representation *C*.

#### c) Reduction of fragment density and rendering

Our goals in reducing fragment density were to obtain a spatially maximally homogeneous representation with fragments placed at positions with highest overlap. To this end, we first sorted the list obtained from step b) according to the quality of overlap in descending order. After that, we adopted an iterative procedure where we first selected the pair of elements *i, j* with smallest distance, and then discarded the element with the smaller overlap with index *k* = max(*i, j*) from the list. This procedure was repeated until a desired fraction *f*_*rem*_ of the original *N*_*rem*_ elements in the list remained. For *f*_*rem*_, we selected nine percentage levels using the function from Torfs et al., 2010:

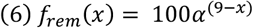

Here, *α* is 0.77 and *x* is the index of the current fragmentation level (*x* = 1, 2, … 9). This yielded the fragment densities of 100%, 77%, 59%, 46%, 35%, 27%, 21%, 16%, and 12%. Finally, the (reduced) representations were rendered by placing the corresponding shapes 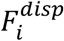 of the curved segment fragments (or of the phosphene) at the selected positions on a black background, scaling the gray levels between 0 (black) and the maximum of the fragments (white).

#### d) Construction of curved segments

For constructing a set of fragments in which not only straight lines, but also corners are well-represented we adopted the approach used in Mineault et al., (2013): each fragment consists of two pointers resembling the hands of a clock. From the fragment’s center, the first hand can point into one of 8 equally spaced directions (0, 45, 90, 135, 180, 225, 270, 315). The second hand can form an angle of 45 (acute angle), 90 (right angle), 135 (obtuse angle), and 180 (straight line) with the first hand. Not counting identical configurations, there are 28 different curved segments which can be constructed by this procedure (see Figure 1B). For rendering these fragments as an image path *F*^*disp*^, we assume a pointer length of *l* = 0.18 arcdeg extending from the center into two directions, and a Gaussian profile with *σ*_*clk*_ = 0.07 arcdeg in dependence on the shortest distance of any point to the two pointers. For obtaining an orientation-resolved representation *F*^*filt*^ of the same fragment, the pointers are rendered separately in the orientation planes which correspond to their direction modulo 180.

### 4. Labelling task

The aim of the labelling task was to identify a unique and unambiguous label for each of the recognizable fragmented objects. Labels are nouns which participants assigned to each of the fragmented objects presented. These labels (i.e., nouns) constitute the set of correct responses for the object recognition task.

#### a) Trial structure and procedure

A fixation point was presented for 500 ms, and then an object was presented until a response was made. The object was rendered with curved segments at a fragment density of 100% (for example stimuli see Figure 2, middle row). The task of the participants was to type the name of the object (a prompt was visible below the object). After entering their response, the participants had to press the key “enter” to start a new trial with a novel object presented. If participants did not recognize the object, they were instructed to type “nan” (for “not a number”).

The participants in the labelling task performed 184 trials (one trial per object). The order of presentation was randomized within participants. To familiarize the participants with the task, they performed four training trials subsequently excluded from analyses.

#### b) Data Analyses

All labels provided by participants were treated as ground truth. Therefore, no performance was extracted from the labelling task. Instead, we assessed the consistency of the labels that participants assigned to a given object. To quantify this consistency, we calculated the label agreement following the methods in Brodeur et al., (2010). For each object, we identified the most frequent label (referred to as the modal name in Brodeur et al., (2010), Brodeur et al., (2012) and Brodeur et al., (2014) after excluding the “nan” responses. We then calculated the percentage of participants who agreed on this label (called modal name agreement in Brodeur et al., (2010), Brodeur et al., (2012) and Brodeur et al., (2014)). The higher this percentage, the greater the label agreement.

We assessed the reliability of the collected labels by comparing them with the ones gathered in (Brodeur et al., 2012). In their study, Canadian French labels from 30 participants were collected for 480 colored object images from the BOSS, including 42 images used in our study. Consequently, both our participants and those in (Brodeur et al., 2012) assigned labels to the 42 objects, albeit in different visual representations. A correlation between the label agreement of both studies was also performed.

To be included in the object recognition task as a “correct response”, a label had to fulfil two criteria. First, as we wanted the labels to reflect a consensus amongst participants, we did not consider objects for which the proportion of “nan” responses exceeded 15%. Second, at least 85% of the participants should agree on the label. Applying these criteria resulted in the exclusion of 35 objects after the first criterion and an additional 105 objects after the second, leaving us with a total of 44 objects. To keep the duration of the object recognition task acceptable for participants, we further refined the selection to include only the 31 objects with highest agreement.

### 5. Object recognition task

To determine the minimum number of fragments necessary to recognize objects, we adapted the object recognition task from Torfs et al., (2010). In this task, fragmented objects are presented with an increasing number of fragments until recognition. We instructed the participants to name the fragmented objects orally (free naming). If they did not recognize the objects, they were instructed to guess.

To avoid participants to see the same object both in phosphenes and curved segment representations, we randomly split the 46 observers into two groups. The first group viewed half of the objects as phosphenes and the other half as curved segments. The second group was presented with the counterbalanced condition.

#### a) Trial structure

We started with a fragmentation level of either 12% phosphenes or 12% curved segments, randomly chosen for each participant. If the response was correct, a novel object was presented at the initial 12% fragmentation level. If the response was incorrect, the same object was presented again, but with the next fragmentation density until the correct object name was given. Up to 9 fragmentation levels could be presented to participants until they gave a correct answer (12%, 16%, 21%, 27%, 35%, 46%, 59%, 77% and 100%). When participants reached the maximum fragmentation level of 100% and still provided an incorrect answer, we showed them the entire object contour (as shown in Figure 1A). This control trial makes sure that a poor recognition of the fragmented object was not due to unfamiliarity with the object. In case a participant was not able to correctly name the full contours of the object, the trials related to the current object were deleted for the current participant (2.22% trials excluded in total). Figure 3 illustrates the time course of the trial.

**Figure 3.**
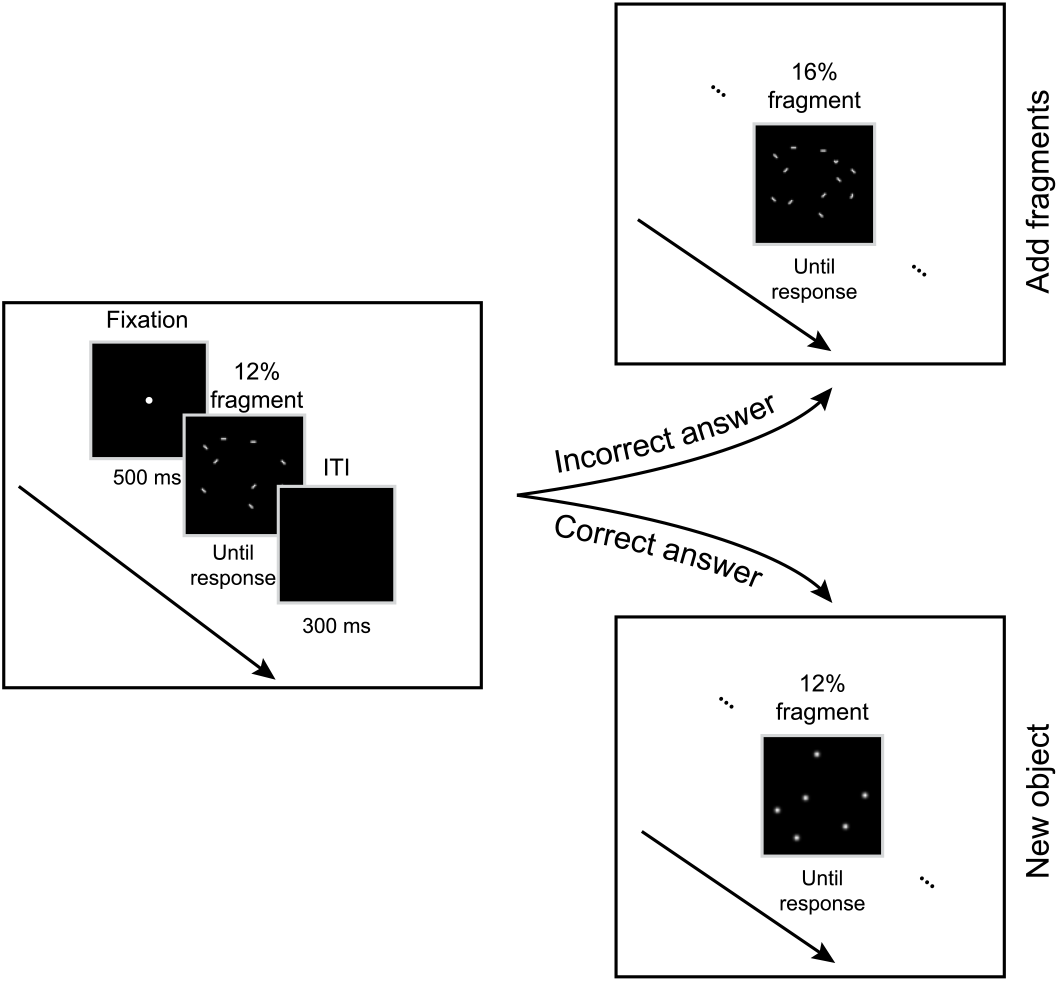
After a fixation point, an object is presented with 12% of either curved segments or phosphenes. The participant is instructed to name the fragmented object out loud. If the response is correct, another object is presented either as phosphenes or as segments with the initial 12% fragmentation level. If the response is incorrect, the same object is presented again with more fragments until the response is correct. There are nine possible fragmentation levels presented in the following order: 12%, 16%, 21%, 27%, 35%, 46%, 59%, 77%, 100%. The trial structure stays the same.

Participants’ answers were rated in real time by an experimenter sitting behind the participants. A response was considered correct either if it matched the label collected in the labelling task, or if it was more specific than the label (e.g., label is “pants” and participants say “jeans”). Otherwise, the response was considered incorrect.

#### b) Procedure

Participants in the object recognition task completed 31 blocks, with each block consisting of up to 10 trials. The number of blocks corresponds to the number of objects selected from the labelling task. The number of trials corresponds to the number of fragmentation levels needed to recognize objects (including full contours). As each participant needed a different fragmentation level to recognize objects, the number of trials performed varied across participants (*M* = 233, *SD* = 17). The order of objects (i.e. blocks) was fully randomized within participants. To familiarize the participants with the task, they performed four training blocks. Objects presented during training were identical for all participants and were not presented during the experimental blocks.

#### c) Data analyses

The performance of participants is determined by the level of fragmentation at which they correctly identified each object (12%, 16%, 21%, 27%, 35%, 46%, 59%, 77% or 100%). The achieved fragmentation level corresponds to the minimum fragment density needed for object recognition. Therefore, the lower the fragment density, the higher the performance. When participants provided an incorrect answer at the 100% level and a correct answer for the full contours of the object, a performance score of 130% was given. This performance score was defined by adding an additional fragmentation level in Equation (6). If some participants failed to recognize the object even when presented with full contours, we discarded the block from analyses for the current participant (2.22% trials excluded in total). Out of the 31 objects analyzed, one was removed for all participants because 15% of participants could not recognize its full contours.

Our hypothesis was that fewer curved segments are needed to recognize objects compared to phosphenes. To test this, we used a linear mixed model to predict the fragment density needed to recognize objects. The type of element (phosphenes vs. curved segments) served as a fixed effect, and individual objects were treated as random effects, allowing both the intercept and slope for each object to vary. We used the following mixed model to predict the needed fragment density for object recognition:

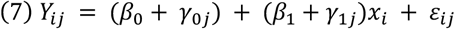

where *β*_0_ is the global intercept (fixed intercept), *γ*_0*j*_ is the intercept of object j (random intercept), *β*_1_ is the slope of fragment type (fixed effect), *γ*_1*j*_ is the random slope of object *j* (random effect) and *ε*_*ij*_ is the residual of object j for a given observation *i*.

We used the R package *MuMIn: Multi-Model Inference* (Bartoń, 2009) to calculate pseudo *R*^2^ for the mixed model. For object-specific analyses, we conducted multiple Mann-Whitney-*U*-tests comparing the number of phosphenes and curved segments needed to recognize each object. We chose a non-parametric approach as the number of observations per object varied due to the inability of some participants to recognize the full contours of some objects. Furthermore, the assumption of homoskedasticity was violated. We applied False Discovery Rate (*FDR*) corrections to adjust the *p* values for multiple comparisons. The effect sizes of each of the Mann-Whitney-*U*-test were calculated with the R package *rstatix*(Kassambara, 2021) providing rank-biserial correlation coefficients (*r*s). An a priori power analysis computed with *G*Power* (Faul et al., 2007) revealed that 42 observations (21 per fragment type) are necessary to detect a significant effect size of *d* = 0.8 (approximately corresponding to *r* = 0.37) with a one-tailed Mann-Whitney-*U*-test (α = 0.05, power = 80%).

## Results

The first aim of our study is to determine the minimum number of phosphenes required for object recognition by rendering only a few phosphenes around the objects’ contours. The second aim is to assess how small changes in the local visual information of phosphenes contribute to object recognition when only a few fragments are present along the objects’ contours. We hypothesized that observers can recognize objects with few phosphenes along the object’s contours. Our second hypothesis states that if observers use the additional amount of local visual information carried by curved segments, they will need less segments to recognize objects compared to phosphenes.

### 1. Reliability of the labels collected in the labelling task

The aim of the labelling task was to identify reliable labels (e.g. nouns) to constitute the set of correct responses for an object recognition task performed by another batch of participants. We selected a subset of images from the Bank of Standardized Stimuli (BOSS; Brodeur et al., 2010) and used an algorithm to represent these images as collections of curved segments at 100% density.

Eighty-one percent of these images (*n* = 34) received identical labels in our study and in Brodeur et al., (2012) where Canadian French speakers saw the unprocessed intact colored images we fragmented. This indicates a high level of label consistency across different participants and viewing conditions. Out of the 42 images considered in our reliability analysis, only 10 had different labels than Brodeur et al., (2012). Four of the labels diverged because our participants were unable to correctly identify the objects presented as curved segments. The remaining six labels differed due to linguistic differences between Canadian and Swiss French (e.g., Canadian French: “suce”; Swiss French: “tétine”).

Additionally, there was a positive correlation (*r* = 0.41, *p* < 0.01, Figure 4) between the label agreement in our study and that in Brodeur et al., (2012). This indicates that a higher agreement between participants of our experiment is associated with similar consensus levels in Brodeur et al., (2012).

**Figure 4.**
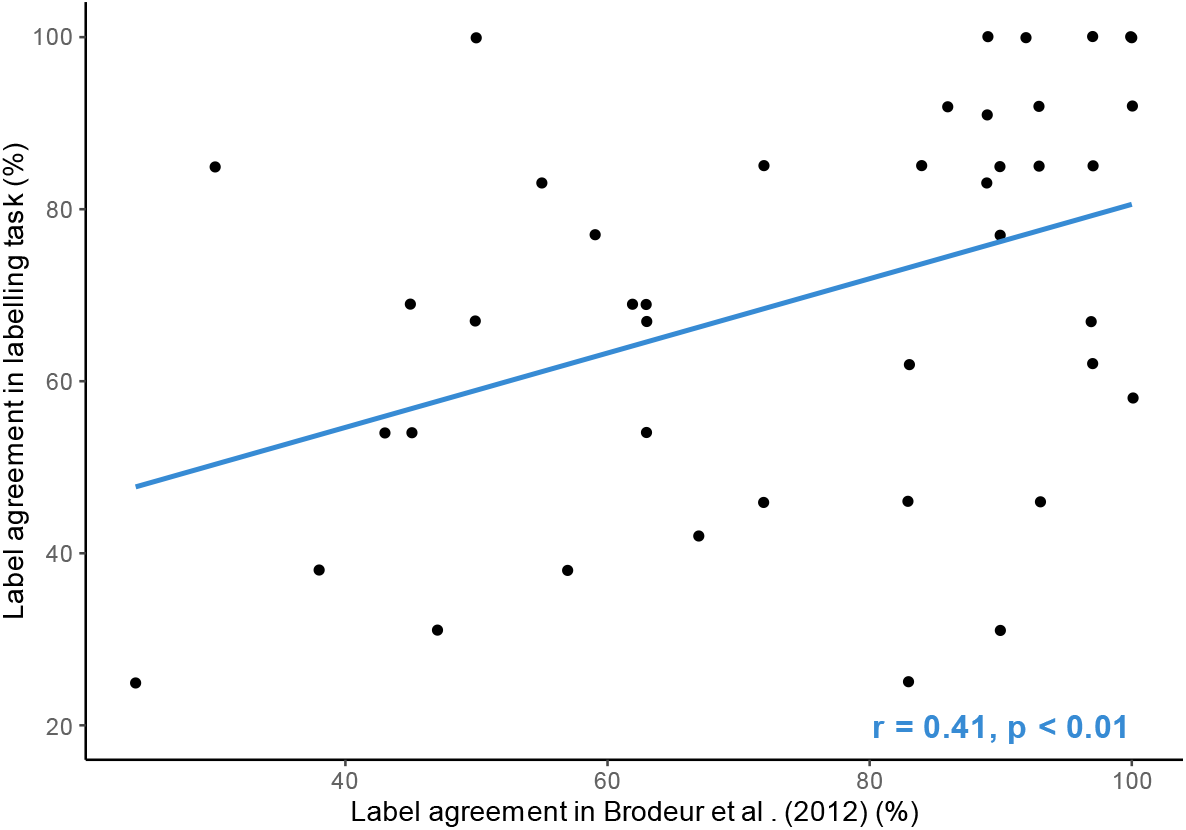
Correlation between the label agreement (in percent) in our labelling task and in Brodeur et al. (2012). The label agreement refers to the percentage of participants who gave the most given label to each object. Each dot on the scatterplot refers to the label agreement of each object presented in both studies (n = 42). Brodeur et al. (2012) presented colored RGB images of objects. In the labelling task, the same objects were presented but fragmented with curved segments with 100% fragment density.

Together, these results suggest that the labels gathered in our study are reliable even with the added complexity of recognizing objects from fragmented images. We conclude that our labelling task generated reliable ground truth for the set of correct responses in a subsequent free naming object recognition task. We selected the 30 objects with highest naming agreement to use as stimuli for our object recognition task. These are shown in Figure 5 along with their corresponding label and label agreement.

**Figure 5.**
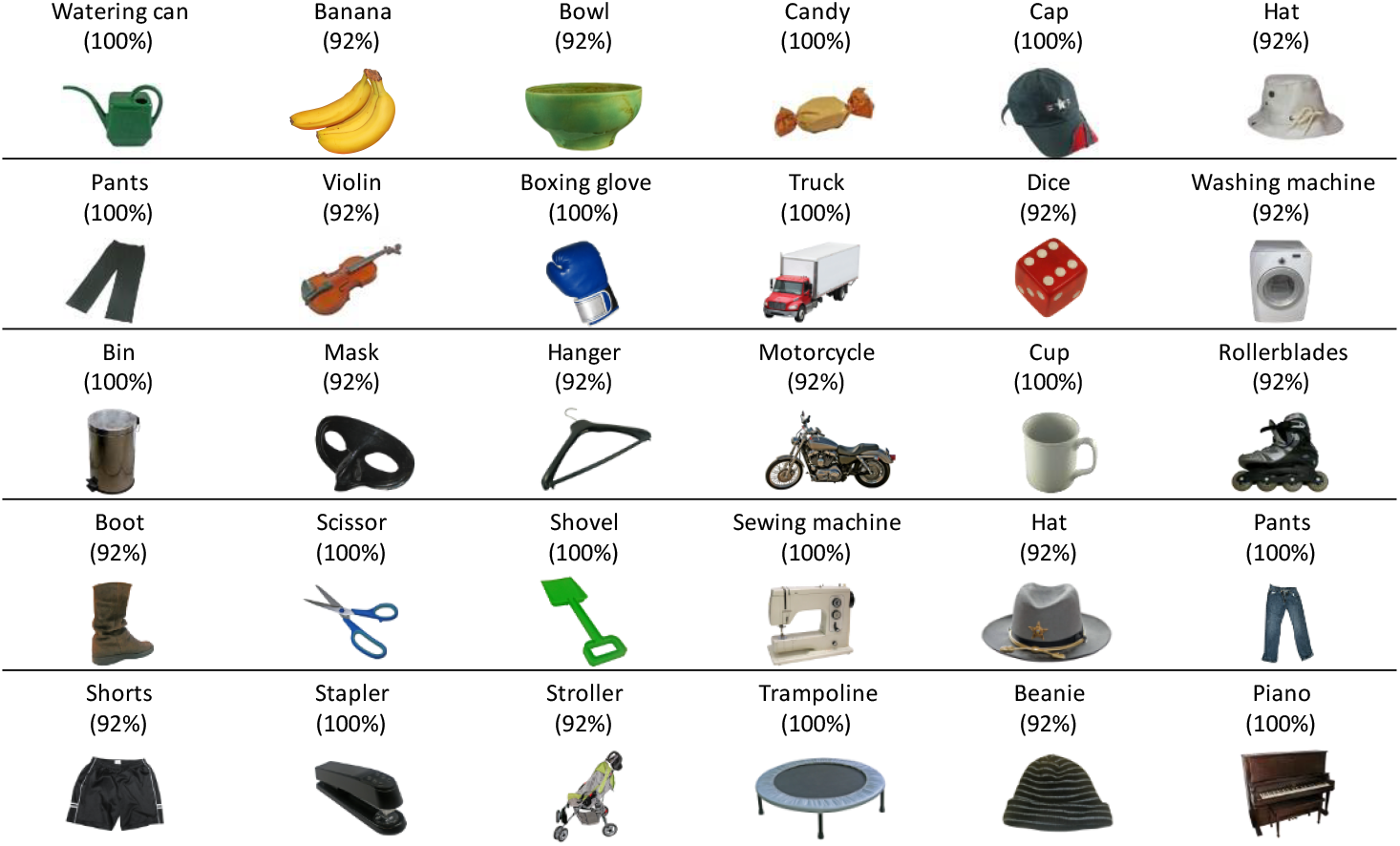
Images selected for the object recognition task along with their translated label (original labels are in French) and the corresponding label agreement. The label is defined as the most widely given name for the current object. The label agreement refers to the percentage of participants agreeing with the label. All colored images were taken from Brodeur et al., (2010) and are licensed CC-BY-NC-SA.

### 2. Contribution of local visual information to object recognition

We hypothesized that observers would need fewer curved segments to recognize objects compared to phosphenes. To investigate the effect of fragment type on object recognition, we conducted a linear mixed model with *fragment type* as fixed effect and individual objects as random intercept and slope. A significant effect of *fragment type* showed that participants needed 27% fewer curved segments to recognize objects compared to phosphenes (β = -27.49, *SE* = 3.48, *p* < 0.001, *R*^2^_fragment type_ = 0.14; Figure 6). This result suggests that observers use the increased amount of local visual information provided by the curved segments to recognize objects.

**Figure 6.**
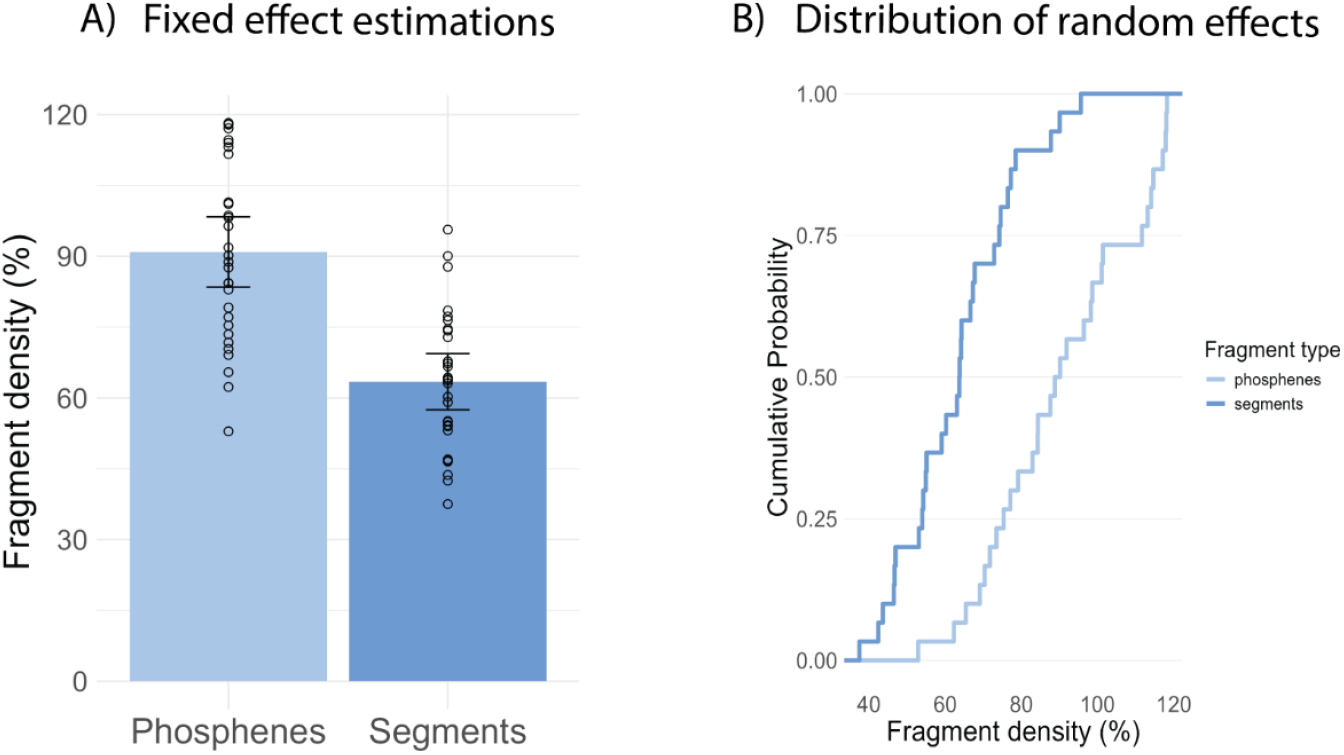
Minimum number of fragments needed to recognize objects. A) Fixed effects estimation from linear mixed model. Bars represent the estimated number of fragments needed to recognize objects when represented as phosphenes and curved segments, respectively. Single dots are estimated random effects of individual objects (n = 30). Error bars are 95% confidence interval. B) Distribution of random effects. Empirical cumulative distributions of estimated random effects of individual objects (n = 30). In both panels, data points above 100% fragment density indicate that objects were not recognized at the maximum fragment density of 100% (performance set as 130%).

The variance explained by the model increased by 23.67% when individual objects were added as random effects (*R*^2^_full model_ = 0.38). This suggest that even though *fragment type* have an effect on object recognition, the minimum number of fragments needed for object recognition still varies depending on the objects. To investigate how strong the effect of *fragment type* varied across objects, we conducted multiple Mann-Whitney-*U*-tests for each object separately. Results showed that 23 out of the 30 objects presented (76.67%) needed significantly less curved segments to be recognized compared to phosphenes (0.001 < *p*s < 0.05; *FDR*-corrected for multiple comparisons). Effect sizes were mid-to high, ranging from *r* = 0.31 to *r* = 0.81 (*r*_mean_= 0.54, *r*_SD_= 0.15), where *r* is the rank-biserial correlation coefficient, a standardized effect size for Mann-Whitney-*U*-tests. Object-specific results are shown in Figure 7. Together, these results show that both the local visual information and higher-level properties such as the object’s type play a role for the recognition of fragmented objects.

**Figure 7.**
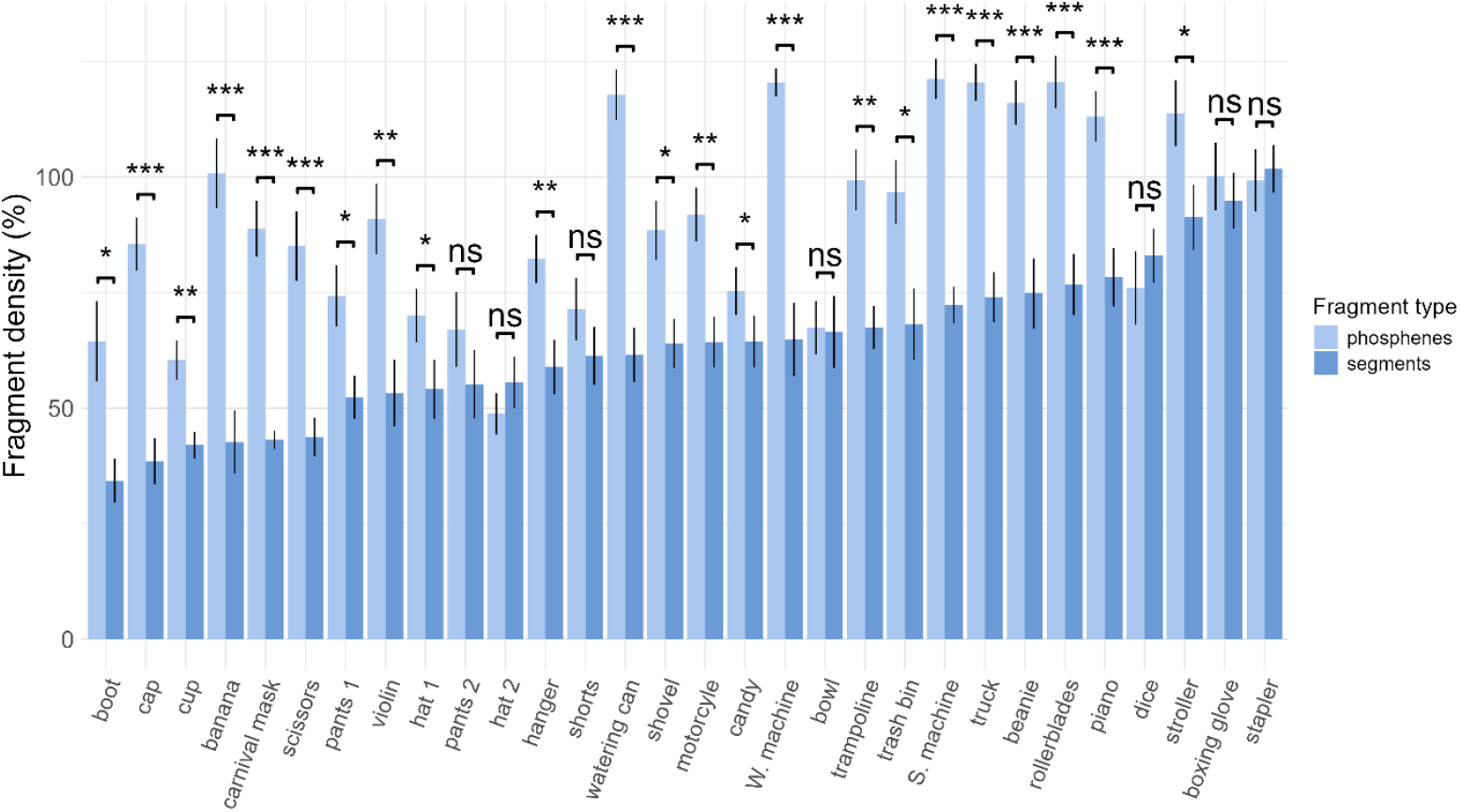
Object-specific effect of fragment type. Each bar is the mean number of phosphenes vs segments needed to recognize each object over all participants. Error bars denote standard errors of the mean. Values above 100% indicate that objects were not recognized at the maximum fragment density of 100% (performance set as 130%). We computed multiple Mann-Whitney-U-tests to assess the difference in the minimum number of phosphenes vs. segments to recognize each object (p < 0.05*, p < 0.01**, p < 0.001***; FDR-corrected for multiple comparisons).

### 3. Minimum number of phosphenes needed for object recognition

To provide a direct proof-of-principle for our minimalistic approach, we back transformed the estimated phosphene density (%) into an estimated number of phosphenes needed to recognize objects. Seventy percent of objects could be recognized when phosphenes were put along their contours. Amongst the recognized objects, participants needed on average 46 phosphenes (*SD* = 10.09) to recognize objects with a minimum of 29 and a maximum of 65 phosphenes. In line with our hypothesis, these results indicate the possibility to recognize objects even very few phosphenes if we sparsely stimulate along the outline of the contours. This is well in line with our minimalistic approach of object rendering in visual prostheses.

## Discussion

Because of the limited resolution of current visual prostheses, smart techniques are required to determine how to best render objects visible with the smallest number of phosphenes possible. As phosphenes are visually isotropic, they provide little-to-no visual information for object recognition. Our second aim was therefore to identify the extent to which small changes in local visual information of phosphenes contribute to object recognition. To answer these questions, we used an adapted study design from Torfs et al., (2010) which gradually increase the number of fragments to quantify the minimum number of curved segments and phosphenes necessary to recognize objects.

We first hypothesized that participants will be able to recognize objects even with few phosphenes placed around objects’ contours. Well in line with this hypothesis, our results showed that, on average, 21 of the 30 presented objects could be recognized with a maximum of 65 phosphenes fitting well the number of phosphenes elicited by current cortical stimulation (Fernández et al., 2021). Some of these objects, for example the hat, could even be recognized with 29 phosphenes. These results contrast with the findings of Zhao et al., (2010) who reported that the threshold for recognizing objects in simulated prosthetic vision was somewhere between 256 and 576 phosphenes. Zhao et al., (2010) tessellated the visual field with various implant resolutions ranging from 8×8 to 64×64, where lower resolutions were simulated with large phosphenes and higher resolutions with small phosphenes. In contrast, we assumed a fixed size for all phosphenes regardless of their density. Therefore, our simulation corresponds the high-resolution cases in Zhao et al., (2010), where we sampled a few phosphenes around the contours instead of covering the entire contours. Our approach shows that in these cases the number of phosphenes can drastically be reduced to less than 100 even for complex objects.

Second, in line with our second hypothesis, we show that approximately 27% less curved segments were necessary to recognize objects compared to phosphenes. In line with Keane (2018), we suggest that participants perform contours’ abstraction to recognize objects when very few curved segments are presented. However, when the segments’ density increases, segments can be interpolated into contours, facilitating object recognition (Persike & Meinhardt, 2017). While the recognition of objects fragmented with curved segments might be supported by these two joint mechanisms, no contour interpolation is possible with the anisotropic local information of phosphenes. We therefore suggest that the recognition of objects fragmented with phosphenes might depend on contours’ abstraction only, explaining why more phosphenes were needed to recognize objects compared to segments. While we did not directly test for disentangling contours abstraction from contours interpolation, one might develop a new experimental design to disentangle these processes.

A closer look on the results showed that the effect size of fragment type on object recognition strongly depended on the object currently presented. Adding the objects as random effects in the linear mixed model showed an increase of 23.67% of explained variance. This suggests that though the local visual information plays a role in object recognition, higher-level information such as the object category is playing a crucial role as well. This suggests that minimizing the number of phosphenes needed for object recognition requires finding an optimal number for each specific object category.

A limitation of our study is that the phosphenes were not rendered in a biological plausible way. We indeed assumed what is informally called a “scoreboard” model in which all phosphenes are rendered homogeneously across the visual field of patients. In the future, one might use a more faithful phosphene representation as the one in (Van Der Grinten et al., 2024). Future studies could also investigate whether the local visual information and the object type interact to understand if observers use different amount of local visual information depending on objects.

## Conclusions

In summary, our study demonstrated that healthy sighted participants could recognize objects with as few as 65 phosphenes sparsely placed around objects’ contours, suggesting that minimalistic rendering could enable object recognition even with the low resolution of current cortical implants. We found that small changes in the local visual information of phosphenes significantly impact object recognition, although this effect varied greatly across different objects. These findings indicate that a homogeneous method for rendering all objects may not be optimal. Instead, an object-specific optimum of phosphene configuration could minimize the number of phosphenes needed for recognition.

## Acknowledgement

Funding was provided by ERA-NET Neuron I-See “Improving intracortical visual prostheses using complex coding and spontaneous activation states” (32NE30_198552; BMBF 01EW2104A) and the Stiftung Bremer Wertpapierboerse.

## Notes

### Competing Interest Statement

The authors have declared no competing interest.

### Summary of Updates

Figure 7 has been revised. The x-axis has been corrected.

